# Full-length tRNAs lacking a functional CCA tail are selectively sorted into the lumen of extracellular vesicles

**DOI:** 10.1101/2024.05.12.593148

**Authors:** Chantal Scheepbouwer, Ernesto Aparicio-Puerta, Cristina Gómez-Martin, Monique A.J. van Eijndhoven, Esther E.E. Drees, Leontien Bosch, Daphne de Jong, Thomas Wurdinger, Josée M. Zijlstra, Michael Hackenberg, Alan Gerber, D. Michiel Pegtel

## Abstract

Small extracellular vesicles (sEVs) are heterogenous lipid membrane particles typically less than 200 nm in size and secreted by most cell types either constitutively or upon activation signals. sEVs isolated from biofluids contain RNAs, including small non-coding RNAs (ncRNAs), that can be either encapsulated within the EV lumen or bound to the EV surface. EV-associated microRNAs (miRNAs) are, despite a relatively low abundance, extensively investigated for their selective incorporation and their role in cell-cell communication. In contrast, the sorting of highly-structured ncRNA species is understudied, mainly due to technical limitations of traditional small RNA sequencing protocols. Here, we adapted ALL-tRNAseq to profile the relative abundance of highly structured and potentially methylated small ncRNA species, including transfer RNAs (tRNAs), small nucleolar RNAs (snoRNAs), and Y RNAs in bulk EV preparations. We determined that full-length tRNAs, typically 75 to 90 nucleotides in length, were the dominant small ncRNA species (>60% of all reads in the 18-120 nucleotides size-range) in all cell culture-derived EVs, as well as in human plasma-derived EV samples, vastly outnumbering 21 nucleotides-long miRNAs. Nearly all EV-associated tRNAs were protected from external RNAse treatment, indicating a location within the EV lumen. Strikingly, the vast majority of luminal-sorted, full-length, nucleobase modification-containing EV-tRNA sequences, harbored a dysfunctional 3’ CCA tail, 1 to 3 nucleotides truncated, rendering them incompetent for amino acid loading. In contrast, in non-EV associated extracellular particle fractions (NVEPs), tRNAs appeared almost exclusively fragmented or ‘nicked’ into tRNA-derived small RNAs (tsRNAs) with lengths between 18 to 35 nucleotides. We propose that in mammalian cells, tRNAs that lack a functional 3’ CCA tail are selectively sorted into EVs and shuttled out of the producing cell, offering a new perspective into the physiological role of secreted EVs and luminal cargo-selection.

## INTRODUCTION

Extracellular vesicles (EVs) are a diverse family of membrane-enclosed structures, actively released from cells into their extracellular environment, including biological fluids^1,2^. EVs are generally categorized into two major size-based subtypes, which are small EVs (sEVs) typically less than 200nm in size, and large EVs (lEVs) exceeding 200 nm in diameter^3^. Interest into EV biology increased since RNA associated with these vesicles^4,5^ may be exchanged between donor and recipient cells^6–8^. In addition, the identification of plasma EV-associated miRNAs that could serve as potential blood-based biomarkers^9–12^ further highlighted the clinical relevance of deciphering the composition of EV-associated RNAs. In that regard, high-throughput sequencing has provided most of the transformative insights into the nucleic acid content of EVs, although most studies focused on short ncRNAs shorter than 50 nucleotides (nt)^5^. Less is however known about the composition and abundance of longer ncRNA molecules associated with EVs^13^. To address this gap, specifically tailored next generation sequencing (NGS) protocols have been applied^14,15^ to mitigate the length bias introduced by the library preparation strategies used in conventional small RNA sequencing protocols^16–20^. Despite these advancements, accurate quantification of highly structured extracellular ncRNAs bearing post-transcriptional modifications, which interfere with reverse transcription^21^, still poses a significant challenge. This difficulty is particularly pronounced for transfer RNAs (tRNAs), which are typically 76-90 nt in length, and may contain up to 13 nucleobase modifications per molecule^22^. In addition to modifications, mature tRNAs are characterized by the presence of a CCA sequence at their 3’ termini, added post-transcriptionally by TRNT1, a tRNA-nucleotidyl transferase that is ubiquitous in eukaryotes^23^. This sequence is crucial for the aminoacylation process, catalyzed by tRNA synthetases^24^, and for the correct positioning of tRNAs within the ribosome^25^.

Mature tRNAs are the most abundant molecules within the cell^26^. Because of their critical functions in mRNA translation, these molecules were thought to be mainly regulated globally, to match demand in protein synthesis. Recent studies have however showed that tRNAs are differentially expressed in cells and tissues^27–29^, or in pathologies such as cancer^30,31^. The emergence of multiple high-resolution approaches tailored to the detection and quantification of tRNAs has also spurred investigations into the biological significance of extracellular tRNA^32^. These studies provided new insights into the small ncRNA composition of EVs^33^ and the identification of nicked, non-EV associated, extracellular tRNAs in biofluids^34^. However, conflicting results on the relative abundance of tRNAs^15,35^, as well as their physical association with EVs^33,36^ have been reported, leaving many open questions as to how tRNAs are associated with EVs, both *in vitro*, in EVs derived from cultured cells, and *in vivo*, in EVs isolated from biofluids^37,38^. In addition, the relative abundance of small nucleolar RNAs (snoRNAs), Y RNAs, and vault RNAs (vtRNAs), compared to tRNAs could be biased due to differences in reverse transcription and amplification efficiencies during library preparation steps^39^.

To address these outstanding questions, improving the precision and accuracy in the quantification and characterization of the full EV-ncRNA repertoire is necessary to unravel the basic mechanisms of extracellular RNA biology and their usefulness as non-invasive biomarkers. Therefore, we adapted ALL-tRNAseq^31^, which was developed for the robust quantification of full-length mature tRNAs, to reduce bias in the global assessment of the small ncRNA repertoire in EVs. Our modified ALL-tRNAseq protocol revealed that the lumen of profiled EVs contains full-length and modified tRNAs that lacked, for the major part, functional 3’ CCA ends. In contrast, the overwhelming majority of cellular tRNAs exist as modified tRNAs with intact 3’ CCA tails. The discrepancy between cellular and EV-associated tRNAs suggests selective incorporation of tRNAs with a compromised 3’ CCA tail into EVs. Because this tail is essential for tRNA aminoacylation, EV-tRNAs are thus, for the most part dysfunctional with respect to protein translation. Since the nucleobase modification profiles of tRNAs in EVs closely resembled those of cellular tRNAs, the EV-sorted tRNAs most likely originated from previously functional cytoplasmic tRNAs rather than via a quality control of newly synthesized, hypomodified tRNAs^22^.

## RESULTS

### Optimizing All-tRNAseq for the detection of longer full-length small ncRNA classes in low input samples

In light of the conflicting reports regarding the small RNA content in EVs^15,19,20,33,35,36,39^, we adapted the ALL-tRNAseq procedure, which was originally developed to quantify full-length and processed tRNAs in cultured cells and tissue samples^31^, to assess the composition of the global extracellular small ncRNA repertoire in EV-enriched and EV-depleted fractions, derived from four different cell lines (i.e., RN, MDA-MB-231, HCT116, and HEK293T). For this purpose, we subjected cell culture media to ultrafiltration (UF) to concentrate particles, followed by size-fractionation with Size Exclusion Chromatography (UF-SEC procedure^40^). This method yields bulk EV fractions either enriched or depleted for EVs. The EV-depleted fractions, or non-vesicular extracellular particle fraction (NVEP)^3^, are enriched in protein-particles, soluble protein and HDL/LDL^41–43^. We then analyzed small RNA expression profiles established using the ALL-tRNAseq protocol on total RNA isolated from cells and collected fractions enriched and depleted for EVs (Figure 1A). We initially evaluated the performance of ALL-tRNAseq without protocol adjustments on low-input EV fractions isolated from lymphoblastoid RN and breast cancer MDA-MB-231 cells. We started with the examination of the most frequently described small ncRNAs, and found that tRNAs were the most abundant transcripts in the size range of 18 to 120 nt, while transcripts in the 18 to 30 nt range, such as miRNAs and Y RNA fragments, were barely detectable (<2%; Supplemental Figure 1A). Based on these observations, we opted to select RNAs of a size >30 nt with an extra size fractionation step before the final PCR amplification to further minimize the contribution of overamplified adapter dimers and ultra-small products in these low input samples^14,38^. The adapted low-input ALL-tRNAseq analysis confirmed a dominance of tRNA molecules across all tested cell lines, accounting for 60-85% of all assigned reads in cells, EVs, and NVEP fractions (Figure 1B).

**Figure 1.**
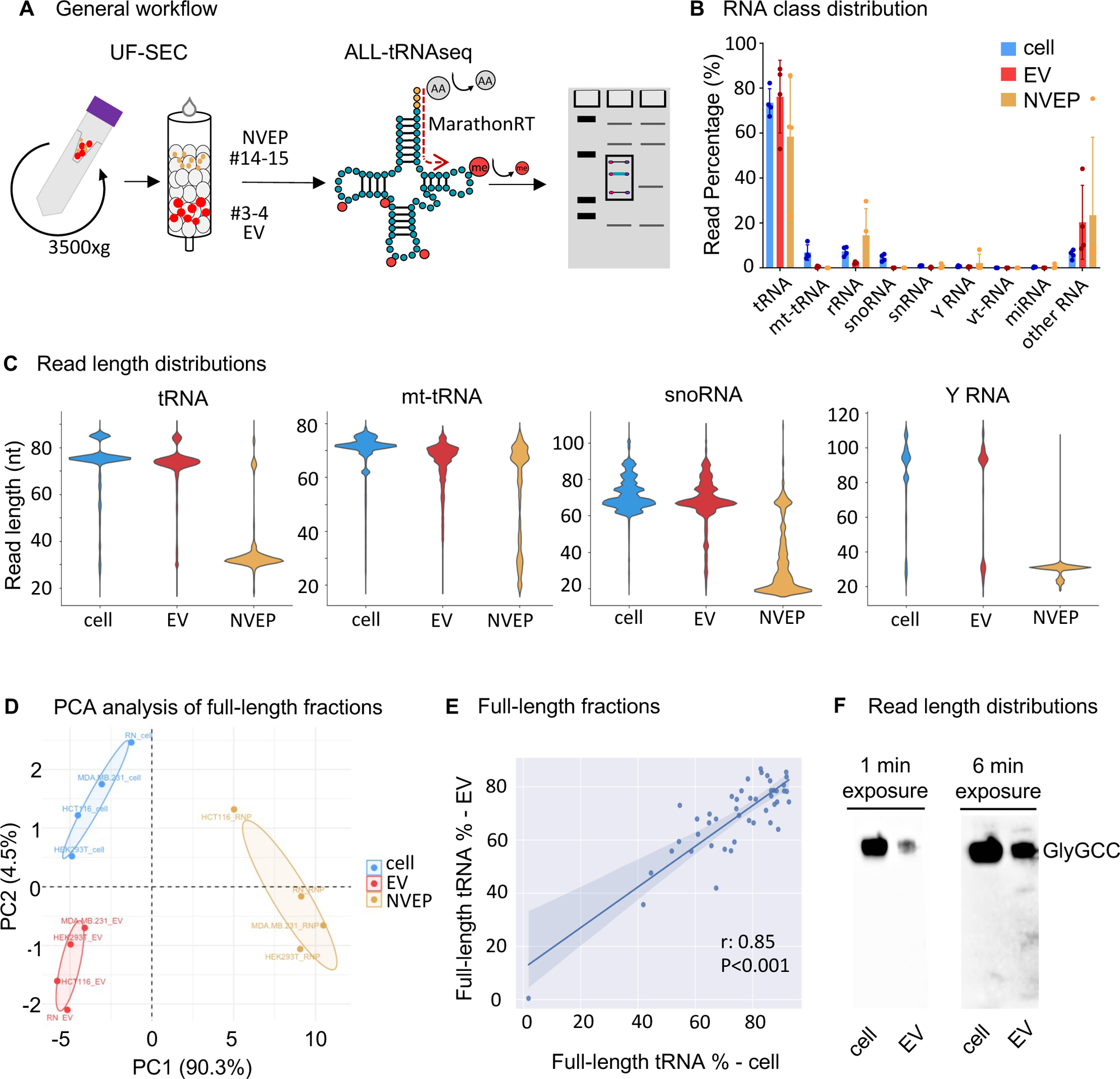
Full-length small ncRNAs are dominating EVs with an overwhelming majority mapping to tRNAs. (A) Schematic overview of the UF-SEC workflow. The collection of EV-enriched and EV-depleted fractions is followed by ALL-tRNAseq on demethylated total RNA isolated from the collected fractions, followed by size-fractionation of PCR amplified sequencing libraries. (B) RNA class distribution in percentage of total normalized reads for the ALL-tRNA-seq library preparation protocol in cells, EVs and NVEP fractions from four cell lines of different origin (RN, MDA-MB-231, HEK293T and HCT116). (C) Violin plots showing read length distributions in percentage of tRNA (first panel), mt-tRNA (second panel), snoRNA (third panel) and Y RNA (fourth panel) in cells, EVs and NVEP fractions from four cell lines of different origin (RN, MDA-MB-231, HEK293T and HCT116). (D) Principal component analysis (PCA) of full-length tRNA anticodon expression in cells (*n* = 4), EVs (*n* = 4), and NVEP fractions (*n* = 4) from 4 cell lines of different origin (RN, MDA-MB-231, HEK293T and HCT116). (E) Pearson correlation (*r*: 0.85) between normalized full-length tRNA reads in cells (n=3; RN) and normalized full-length tRNA reads in EVs (n=3; RN). (F) Detection of tRNA-GlyGCC in cells (left lane) and EVs (right lane) by Northern blot, 1-minute exposure shown in the left blot and 6-minute exposure in right blot.

Early studies on the composition of EV-RNAs have suggested that the EV-associated tRNAs were predominantly shorter than 50 nt^44^. However, more recent studies relying on the use of highly processive group II intron reverse transcriptases, such as TGIRT^15,33,38^, or bioanalyzer assays^45^ reported tRNA lengths of 60-72 nt. We examined the distribution of extracellular small ncRNA lengths using our adapted ALL-tRNAseq, focusing on the most abundant small ncRNA subtypes already described in EVs that encompass full-length transcripts shorter than 120 nt. We could detect full-length molecules for all previously described vesicular ncRNAs (except for miRNAs and rRNAs, which fall, respectively, below and above the size selected range) (Figure 1C). The vast majority of reads mapping to cytoplasmic and mitochondrial tRNAs (mt-tRNAs) appeared to correspond to full-length species, defined here as reads encompassing the complete tRNA sequence encoded in the genome, in both cells and EVs (Figure 1C; first and second panel). Reads mapping to full-length cytoplasmic tRNAs included type I tRNAs (∼75 nt in length) and type II tRNAs (∼85 nt in length), which differ in length due to length differences of the variable loop. Other ncRNAs however, like Y RNAs (RNY1, 3, 4 and 5) and snoRNAs, displayed a wider range of sizes and thus fragmentation or processing (Figure 1C; third and fourth panel). In striking contrast, the size distribution analyses performed in the EV-depleted NVEP fractions revealed that most small ncRNAs within the 18-120 nt size range, existed almost exclusively as fragmented and/or nicked RNAs^34^, with the exception of a modest, but near-intact population of mt-tRNAs (Figure 1C; compare first, third and fourth panels to second panel). Altogether, this data indicated that the most abundant ncRNAs share remarkably similar length distributions in cells and their EVs, while the NVEP fractions demonstrated markedly distinct size distributions in comparison to both cells and EVs (Figure 1C).

### Full-length tRNA composition is highly similar between cells and EVs

Considering the striking dominance of full-length type I and II tRNAs in the established read length distribution profiles (Figure 1C; left panel), we wondered whether variations in tRNA lengths were occurring at the anticodon level. We thus compared the full-length tRNA fractions for each tRNA anticodon in cells, EVs, and NVEP fractions. The results revealed a complete separation between cells and EVs with the NVEP fractions across all tested cell lines (Figure 1D), consistent with the global tRNA length distribution in these samples (Figure 1C). We further analyzed the relationship between the full-length presence of all tRNA anticodons in cells and EVs. This analysis confirmed a robust correlation between the relative abundance of full-length tRNAs across all tRNA anticodons in cells and EVs (Pearson correlation *r*: 0.85; Figure 1E), which were comparable to the ratios observed in samples derived from the other profiled cell lines (*r*: 0.78 to 0.84; Supplemental Figures 1B to 1D).

Next, we aimed to validate our reverse transcription-based sequencing results, particularly the strikingly low abundance of tRNA derived small RNAs (tsRNAs) in EVs, using a hybridization-based approach to assess RNA size distribution. For this purpose, we selected tRNA-GlyGCC, an extensively characterized and abundant extracellular tsRNA identified in cell culture media and biofluids^45–47^. Our results confirmed an overwhelming predominance of full-length tRNAs in both cells and EVs, as similarly observed by Tosar et al^47^. Smaller tsRNAs or nicked tRNAs were only faintly noticeable after an extended period of exposure (Figure 1F). Collectively, these data suggested that EVs are associated primarily with full-length tRNAs that mirrored the composition of the tRNA repertoire in the producing cells.

### The lumen of EVs contains fully processed and modified mature tRNAs

To determine whether the detected extracellular full-length tRNAs are located within the lumen of EVs or bound on the external surface of EVs, we treated EVs with proteinase K, followed by an RNase sensitivity assay. In these conditions, enclosed RNAs are protected from RNAse, while surface-bound RNAs, even if originally bound by proteins, are degraded by the treatment (Figure 2A)^33,34,48^. The RNA sensitivity assay revealed that EV-tRNAs were almost completely protected from the protease and RNase I treatment, reflected by the near-identical length distribution profiles between untreated and treated EV fractions (Figure 2B). We also confirmed that tRNAs were not intrinsically resistant to RNAse I using size-selected cellular tRNAs (Supplemental Figure 1E).

**Figure 2.**
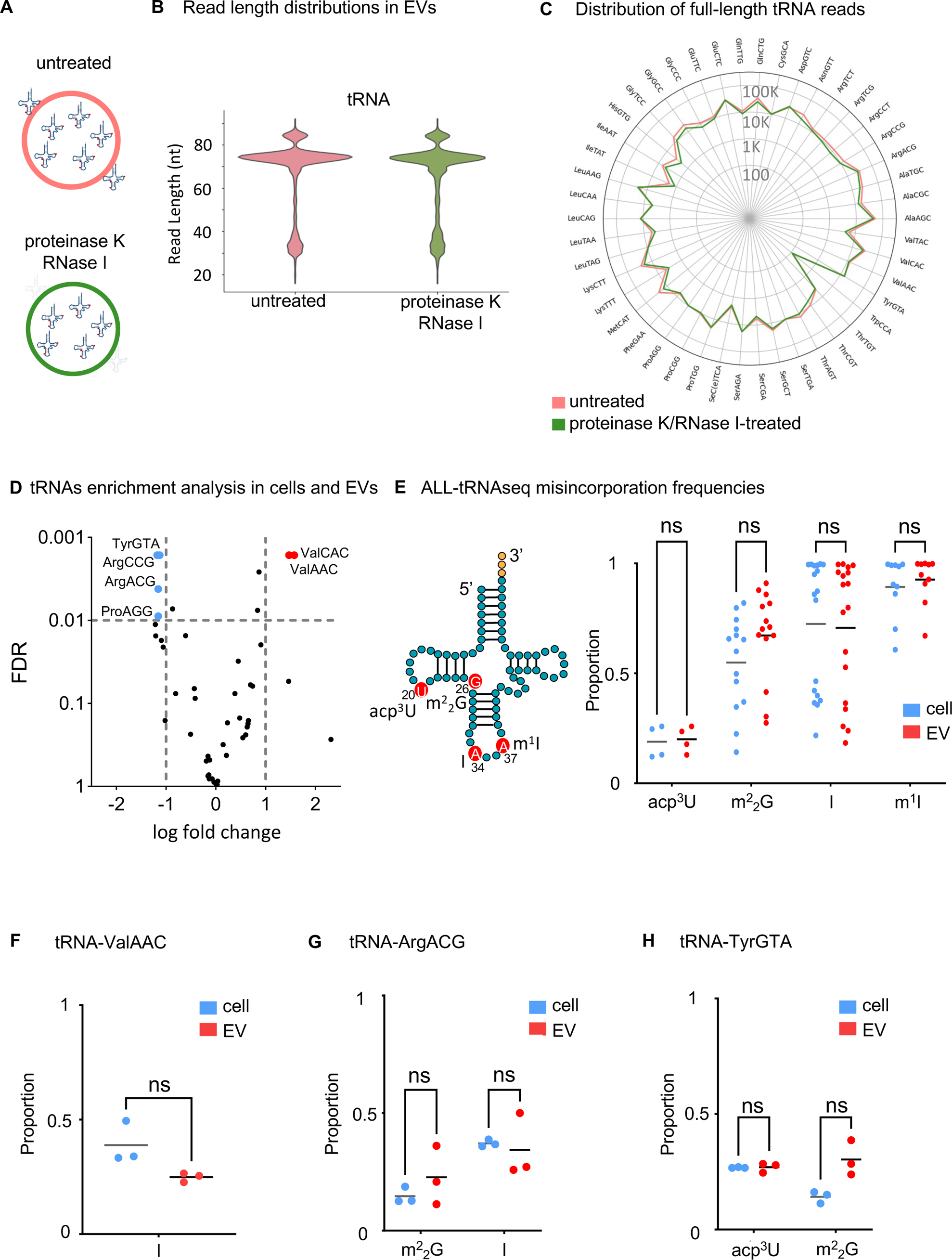
The lumen of EVs contains fully processed and modified mature tRNAs. (A) Schematic representation of the possible localization of EV-associated tRNAs in untreated conditions (top; pink), and upon treatment with proteinase K and RNase I (bottom; green). (B) Violin plots showing read length distribution of total normalized read counts before (left; pink) and after (right; green) treatment with proteinase K and RNase I. (C) Radar plot of full-length tRNA anticodon reads per million mapping to mature cytoplasmic tRNA, showing the read distribution per tRNA anticodon for untreated EVs (red line) and proteinase/RNase I-treated EVs (gray line). Data are represented as Log 10 values on the radius. (D) Volcano plot of significantly differentially expressed tRNAs between cells (n=3) and EVs (n=3). tRNA anticodons were classified according to their fold-change (x axis) and plotted against their FDR values (y axis). Differentially expressed EV-tRNAs are highlighted in red and differentially expressed cellular tRNAs in blue (FDR<0.01; gray line). The vertical gray line represents a 2-fold change. (E) Proportion of nucleotide variations in EV-enriched tRNA-ValAAC from cells (n=3; blue) and EVs (n=3; red). (F) Proportion of nucleotide variations in cell-enriched tRNA-ArgACG from cells (n=3; blue) and EVs (n=3; red). G) Proportion of nucleotide variations in tRNA-TyrGTA in cells (n=3; blue) and EVs (n=3; red). (H) Mismatch analysis showing misincorporation rates T>C/G, G>A, A>G, indicative of acp^3^U, m^2^_2_G, I, and m^1^I at nucleotide positions 20, 26, 34, and 37, respectively. Modification sites are indicated in tRNA secondary structure. All tRNA reads were examined for the presence of mismatches in cells (blue) and EVs (red). Significance for 2E-H was calculated using a Student’s t-test.

For a more comprehensive comparison of tRNA expression between cells and EVs, we examined full-length tRNA expression untreated EVs and proteinase K/RNase I-treated EVs from the lymphoblastoid RN cell line (B cell lymphoblasts that are widely studied for sEV production)^7,49,50^. These results indicated that the abundance of full-length tRNAs was relatively consistent across untreated EVs and proteinase K/RNase I-treated EVs (Figure 2C). To identify whether specific tRNAs were significantly enriched in cells or EVs, we categorized all tRNAs based on their fold enrichment in EVs relative to cellular tRNA reads. This analysis revealed a significant enrichment of tRNA-ValAAC and tRNA-ValCAC in EVs (false discovery rate FDR <0.01), while tRNA-TyrGTA, tRNA-ArgACG, tRNA-ArgCCG and tRNA-ProAGG were more abundant in cells rather than in EVs (FDR <0.01; Figure 2D).

The significant enrichment of valine tRNAs in EVs prompted us to examine whether distinct modification profiles could be detected in EVs, potentially associated with the secretion of these tRNAs via EVs. The precision and consistency of the reverse transcription process are depending on the reaction conditions and the inherent characteristics of the enzyme^51^. Thus, our analysis was limited to misincorporation signatures linked specifically to the use of MarathonRT, excluding methylated residues that are removed with AlkB (m^1^A, m^3^C, m^1^G, and m^2^_2_G) treatment, which is required for the processive reverse transcription of full-length tRNAs^52^. The misincorporation signatures included 3-amino-3-carboxypropyluridine (acp^3^U, position 20), N2, N2-dimethylguanosine (m^2^ G, position 26), inosine (I, position 34), and 1-methylinosine (m^1^I, position 37; Figure 2E), which were previously identified as detectable modification sites for MarathonRT in an independent benchmarking study on tRNA-seq quantification^53^. These signatures were further validated through annotations in the MODOMICs database^54^, and the gtRNAdb database^55^, and were independently confirmed as detectable modification sites for high-throughput sequencing^56–58^. First, we used the overall mismatch profiles unique to ALL-tRNAseq to examine whether the general nucleobase modification patterns in EVs compared to their cellular counterparts were differentially modified regardless of tRNA abundance. When we compared the misincorporation signatures in all tRNA sequences between cells and EVs, we observed minimal differences in overall modification levels (Figure 2E). Given that modifications within the anticodon loop are typically regarded as facilitators of mRNA decoding efficiency and fidelity^59^, we next focused on evaluating tRNAs harboring inosine at position 34, resulting in an A to G conversion. When applied to the significantly EV-enriched tRNA-ValAAC transcripts, this analysis revealed an average of 25% A>G mismatches at position 34 compared to its canonical A, in contrast to an average of 39% in cells (Figure 2F; p=0.06). Likewise, we analyzed m^2^ G (position 26) and I (position 34) for the cell-enriched tRNA-ArgACG, which revealed highly similar modification profiles in cells and EVs (Figure 2G). Given the cell-enrichment of multiple tRNA isoacceptors, we also evaluated another cell-enriched tRNA-TyrGTA, expanding our modification analysis to acp^3^U (position 20) and m^2^ G (position 26) sites. Again, this analysis revealed similar levels of misincorporations at position 20 (T>C or T>G, corresponding to acp^3^U; ns, p:0.9) and 26 (G>A, corresponding to m^2^ G; ns, p: 0.03) in EVs compared to cells (Figure 2H). Additionally, we analyzed tRNA-AlaAGC that contained all detectable modifications sensitive to the detection with MarathonRT, but was neither enriched in cells nor in EVs. Similar to previous results, no statistically significant differences between the misincorporation signatures were observed for tRNA-AlaAGC in EVs (Supplemental Figure 1F). Altogether, these findings suggested that the detected misincorporation signatures were highly similar between cells and EVs, and detected differences in modification levels did not result in differential secretion into EVs. This implies that the tRNAs found in EVs likely originated from the mature full-length tRNA cellular pool, rather than defective or hypomodified tRNAs that are known to be targeted for elimination by a quality control pathway^60^. Overall, these observations indicated that the tRNAs found in EVs were sorted into the lumen of EVs most likely from the mature tRNA population, containing a range of chemical modifications comparable to the cellular tRNA pool.

### Mature full-length tRNAs without a functional 3’ CCA tail are sorted into EVs

Following their transcription, pre-tRNAs are processed into mature tRNAs, which involves removal of their leader and trailer sequences (as well as intronic sequences for the few tRNA genes that harbor an intron), nucleobase modifications, and the addition of a CCA tail at the 3’ end of the tRNA^61^. An intact 3’ CCA tail is essential for tRNA functions in translation as it constitutes the point of linkage for the corresponding amino acid, a process catalyzed by aminoacyl-tRNA synthetases. Interestingly, a size distribution analysis of the full-length tRNA portion between 60-90 nt, which includes type I and type II tRNAs, revealed a broader set of tRNA peaks spanning a size range of 72-76 nt in EVs, as opposed to the expected 75-76 nt size range typically observed for type I tRNAs in cells (Figure 3A). A similar pattern was also evident for the expected ∼85 nt-long type II tRNAs, which appeared to be divided in a series of peaks shortened by up to 3 nucleotides (Figure 3A). To further investigate this intriguing length difference in EV-tRNAs, we analyzed the tRNA reads alignments of a few selected species. To our surprise, we observed a gradual truncation of the tRNA 3’ CCA tail in EVs, which, to a much lesser degree, extended into the mature tRNA sequence (Figure 3B). As this did not seem to occur at the 5’ end of EV-enriched tRNA-ValCAC-1-1 (Figure 3B), and was generally absent in EVs (Supplemental Figure 1G), the observed truncation appeared specific for the 3’ CCA in EVs. This remarkable truncation of the 3’ CCA tail was not restricted to a few tRNA species in EVs derived from RN cells, but was also observed for tRNAs in EVs from MDA-MB-231, HCT116 and HEK293T cells (Figure 3C). In contrast, the vast majority of cellular tRNAs harbor, as expected, an intact 3’ CCA tail (Figure 3C; p<0.001). tRNAs lacking a complete CCA ending displayed an almost uniform distribution of the possible sequences between the intact and complete lack of the CCA sequence (i.e. CCA, CC-, C-- or ---; Figure 3C), which is compatible with the action of 3’ to 5’ exonuclease(s) rather than a precise endonucleolytic removal of the 3’ CCA tail. To further validate these findings, we performed ligation-based quantitative PCR (qPCR) which can distinguish tRNAs with a complete CCA sequence from species lacking this terminal sequence. These assays confirmed a near complete absence of tRNAs lacking the CCA sequence in cells (Supplemental Figure 1H) with only unspecific amplification generated after 40 PCR cycles, as verified on a 6% polyacrylamide gel (Supplemental Figure 1I). In contrast, both CCA-less and CCA-containing tRNAs were readily detected in EVs (Supplemental Figure 1H and Supplemental Figure 1I), confirming the existence of a large proportion of full-length tRNAs lacking an intact 3’ CCA tail in EVs. Lastly, we evaluated the distribution of the CCA truncations in RN cells and EVs across all tRNA isoacceptors. This analysis revealed that 98% of cellular tRNAs contained the crucial CCA sequence for aminoacylation (Figure 3D; top panels), whereas the extent of the CCA truncation (>52%) was similar in all EV tRNAs (Figure 3D; bottom panels). The consistency of this CCA truncation pattern across all tRNA types inside EVs raises the intriguing possibility of a potential sorting mechanism favoring fully processed mature tRNAs that are stripped of their essential 3’ CCA ending.

**Figure 3.**
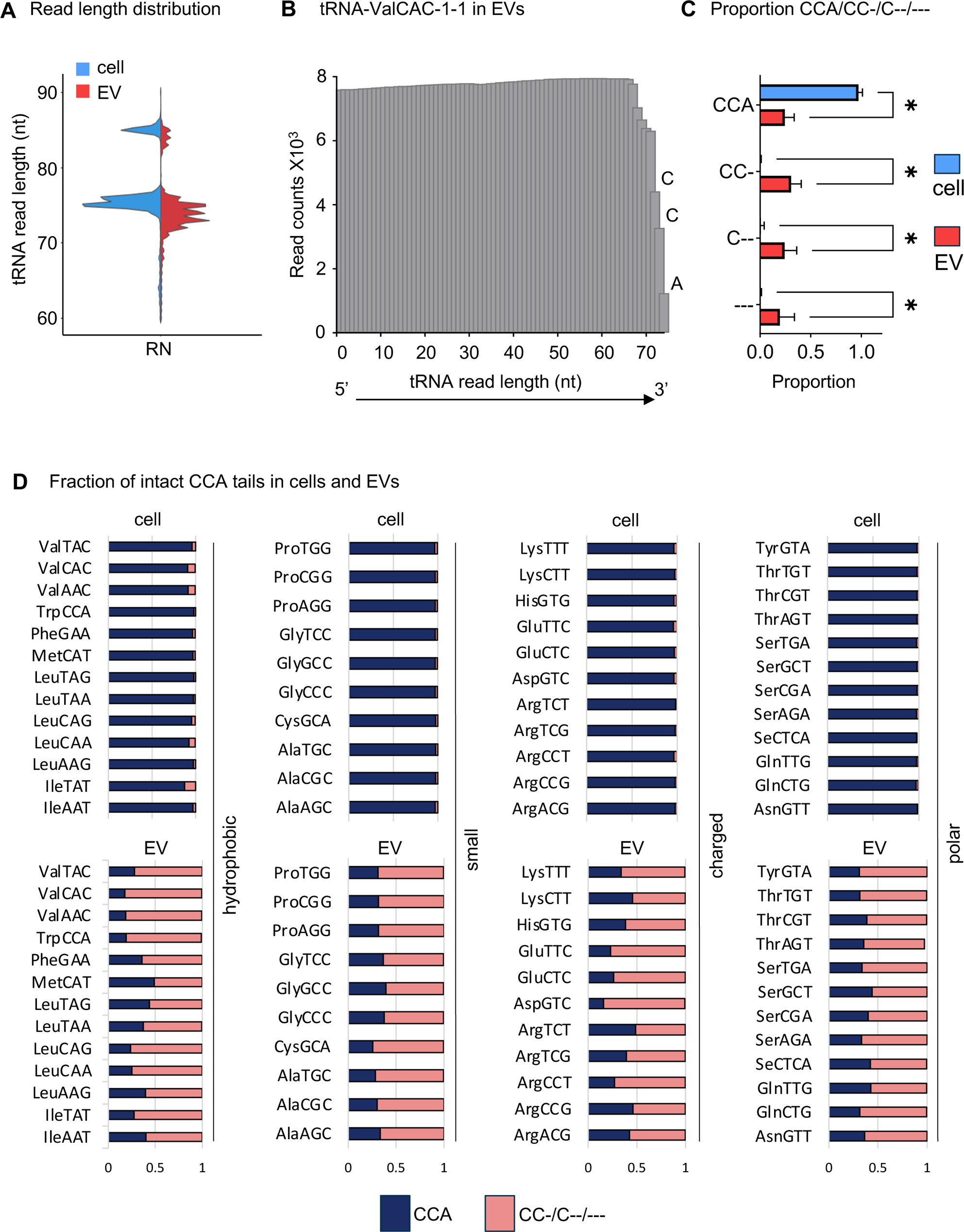
Mature full-length tRNAs without a functional CCA tail are sorted into EVs. (A) Split violin plot showing tRNA read length distribution of 60-90 nucleotides in percentage in cells (left) and EVs (right). (B) Alignment plot showing the cumulative reads for each position along the sequence of EV-enriched tRNA-ValCAC-1-1 in a RN cell line. The x-axis follows the tRNA sequence numbering from positions 1 at the 5’ terminus to 74 at the 3’ terminus. (C) Comparison between cells and EVs in the proportion of tRNA reads containing a CCA/ CC-/ C--/ --- (p<0.001) endings from 4 cell lines of different origin (RN, MDA-MB-231, HEK293T and HCT116). Statistics were performed using unpaired t-tests corrected for multiple testing with False Discovery Rate (FDR). Data represent the average (+/− SD). (D) Fraction of intact CCA endings in cells (n=3; top panels) and EVs (n=3; bottom panels), represented as horizontal bar graphs with a proportion of zero to one. Each horizontal line represents a tRNA type, grouped by anticodon, displaying 46 anticodons in total, which are grouped based on the chemical properties of their cognate amino acids. An intact CCA-tail is indicated in blue, damaged CCA endings are indicated in pink. tRNA anticodon reads are normalized per million.

### Full-length small ncRNA profiles can be recapitulated in plasma EVs

To determine whether EVs derived from a physiologically relevant body fluid are also enriched in CCA-truncated tRNAs, we analyzed the size distribution of EV-associated tRNA in plasma EVs (pEV) from patients with classic Hodgkin Lymphoma (cHL). The sequencing of pEVs using an alternative group II intron RT, known as TGIRT, previously revealed the presence of both full-length and fragmented small RNAs in human plasma samples^38^. Consequently, we analyzed all RNA species in the range of 18-120 nt in our pEV samples. EVs obtained from biofluids are expected to display a much more complex diversity of small RNA molecules compared to cell culture-derived EVs as they originate from a large number of diverse cell types^62^.

Within this size range, cytoplasmic tRNAs remained the predominant small RNA class in pEVs, representing an average of 63% of all mapped ncRNA reads, followed by mitochondrial tRNAs (26%). In contrast, snoRNAs (<1%), snRNAs (<1%), vault RNAs (<1%), and miRNAs (2%) contributed minimally to the RNA pool within the size range analyzed, and were therefore omitted from further analysis (Figure 4A). Previous studies have indicated that full-length Y RNAs are stably associated to EVs in circulation and are present at high levels^48^. Our analysis in pEVs confirmed that Y RNA (8%) indeed displayed a greater prevalence in the circulating pool compared to the analyzed culture media (compare Figure 1B to Figure 4A). Therefore, we next determined Y RNA size distributions in pEVs in more detail, which showed clear differences in the relative abundance of full-length Y RNAs compared to the read distribution profiles observed in cell culture-derived EVs (Figure 4B). Although we did observe full-length Y RNAs in pEV samples, a larger proportion of Y RNA fragments of approximately 30 nucleotides could be appreciated in pEVs (Figure 4B). Additionally, we could observe differences in ratios for full-length Y RNAs, specifically for 96 and 102 nt-long species in pEVs compared to cell culture-derived EVs, corresponding to RNY4 and RNY3 respectively (Figure 4B). Consequently, we performed a dedicated expression analysis of all four full-length Y RNAs in pEVs and EVs derived from cell culture media. Consistent with the Y RNA read length profiles (Figure 4B), this analysis demonstrated that full-length RNY3 and RNY4 are both abundant in pEVs. However, in EVs from cell culture media, full-length RNY4 dominated the total full-length Y RNA read counts, albeit with large variation among the tested cell lines (compare Supplemental Figure 1J to Figure 4C). This data indicated that both the composition of small RNAs and the expression and fragmentation of Y RNAs in circulating pEVs differed from those observed in EVs from cell culture media.

**Figure 4.**
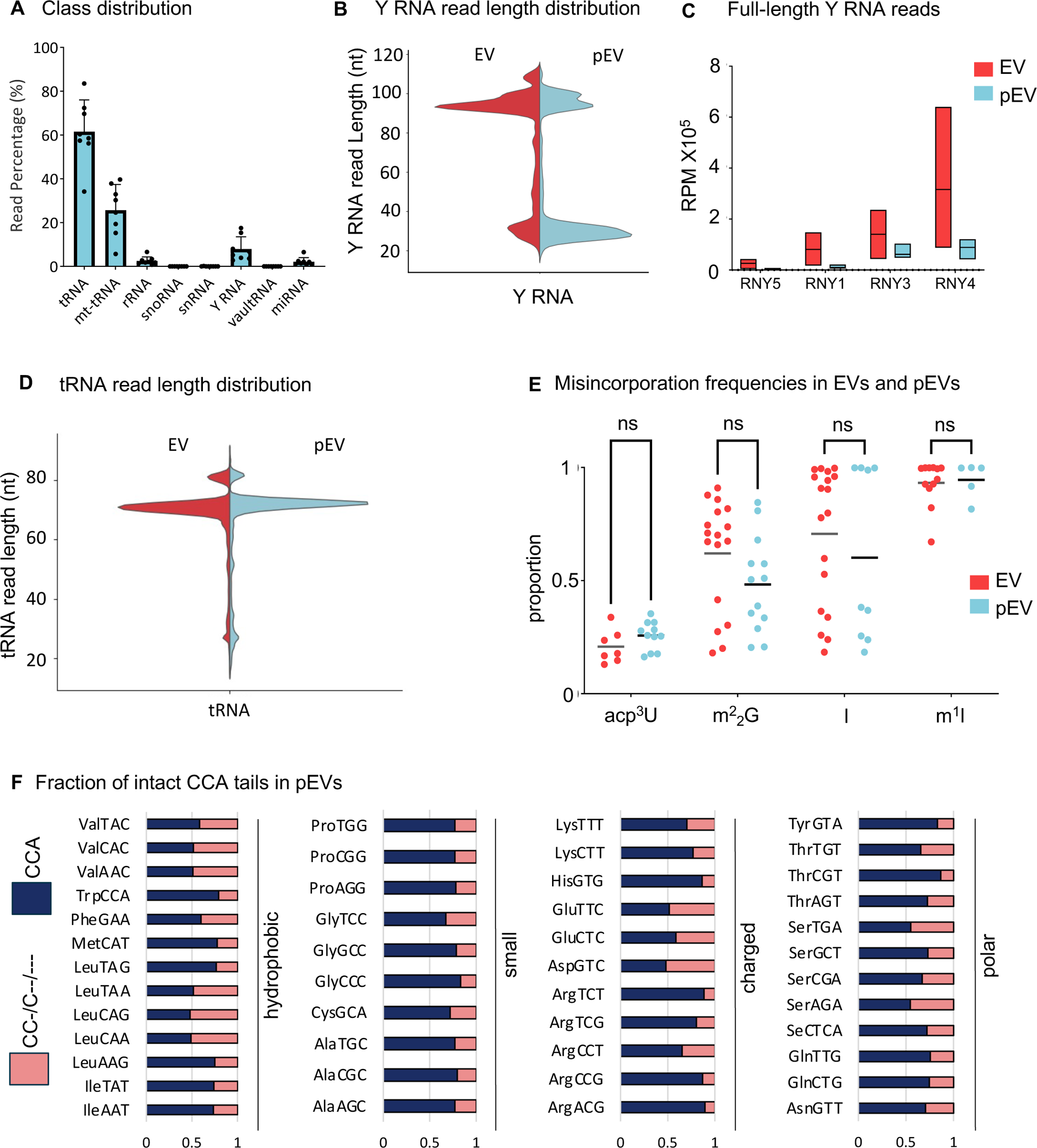
Full-length small ncRNA profiles can be recapitulated in pEVs. (A) RNA class distribution in percentage of total normalized reads for the ALL-tRNA-seq library preparation protocol in pEVs. (B) Split violin plot showing the average Y RNA read length distribution in percentage in 4 EV samples (red) and 8 pEV samples (blue). (C) Expression of Y RNA reads per million mapping to a Y RNA library for EVs and 8 pEV samples. Red dots represent EV samples from culture supernatants, blue dots are pEV samples. (D) Split violin plot showing tRNA read length distribution in percentage in in 4 EV samples (red) and 8 pEV samples (blue). (E) Mismatch analysis showing misincorporation rates T>C/G, G>A, A>G, indicative of acp^3^U, m^2^ G, I, and m^1^I at nucleotide positions 20, 26, 34, and 37, respectively. All tRNA reads were scrutinized for the presence of mismatches in EVs (red) and pEVs (blue). (F) Horizontal bar graphs showing the proportion CCA from 0-100% in cells and EVs. An intact CCA-tail is indicated in blue, damaged CCA endings are indicated in pink. tRNA reads are normalized per million. Each horizontal line represents a tRNA type, grouped by anticodon, displaying 46 anticodons in total, which are grouped based on the chemical properties of their cognate amino acids.

While Y RNAs exhibited high read counts for both full-length and fragmented molecules (Figure 4B), the vast majority of tRNA molecules existed as full-length species in pEVs, similar to the relative full-length tRNA abundance in cell culture-derived EVs (Figure 4D). Next, we compared the misincorporation signatures for acp^3^U, m^2^_2_G, I, and m^1^I in pEVs and EVs collected from cell culture media. This analysis also revealed highly similar modification profiles in pEVs compared to cultured cells and their EVs (compare Figure 2E to Figure 4E). Finally, we assessed the integrity of the 3’ CCA termini in pEVs. While we observed a clear contrast in the presence of intact 3’ CCA tails between cellular tRNAs (ranging from 88 to 98%; Figure 3D, top panels) and cell culture-derived EVs (ranging from 16 to 49%; Figure 3D, bottom panels), the range observed in pEVs spanned from 48 to 89% across all samples (Figure 4F). Altogether, our results indicate that while there are subtle variations in the small ncRNA composition between pEVs and EVs obtained from cell culture media, tRNAs are consistently the predominant component of EVs both *in vitro* and *in vivo* in the 18-120 nt size range. Remarkably, the tRNA profiles in plasma EVs contained comparable modification profiles and included comparable levels of truncated 3’ CCA tails, albeit to a lesser degree than what is observed in EVs collected *in vitro*.

## DISCUSSION

The intricate composition of extracellular small RNAs and the mechanisms governing their sorting into EVs have often been overlooked due to limitations inherent to conventional sequencing-based methods. These methods are hindered by various technical biases, particularly when dealing with low-input samples. In addition, analyses of extracellular ncRNAs have traditionally focused on the class of miRNAs and as a consequence much attention has been given to RNA species or fragments within this size range^18,39^. However, the development of optimized methods for quantifying longer full-length ncRNAs, beyond the miRNA range, has been relatively scarce^15,33,38,46^. Specifically, the detection and quantification of tRNAs pose significant challenges due to the rigid tertiary structure of tRNAs and the presence of highly modified nucleotides in the Watson-Crick face of tRNA bases. Together, these properties may lead to either an RT arrest or nucleotide misincorporations^52^.

The introduction of thermostable group II intron sequencing (TGIRT-seq) suggested the presence full-length tRNAs in EVs^33^. This confirmed observations by Northern blot indicating several individual full-length tRNAs are associated with EVs^47^, but whether this was the case for all EV-tRNAs remained unknown. Whereas protocols using highly processive group II intron RTs indicate a high percentage of tRNAs within EVs, these enzymes fail to overcome modifications at the 5’ end, leading to a large portion of incomplete tRNA sequences predominantly starting after nucleotide U16 of tRNAs^33^. Remarkably, tRNA length profiles from all recently developed methods for low-input samples reported tRNA sizes up to 80 nucleotides. This suggests that type II tRNAs (with expected sizes of 84-85 nt) are neither efficiently nor accurately covered in these datasets^15,33^. Consequently, most prior investigations into the function and biomarker potential of tRNAs in both cell culture and biofluids were limited by standardized sequencing protocols, resulting in a disproportionate emphasis on tsRNAs and an underrepresentation of full-length tRNAs. In the present study, we refined the ALL-tRNAseq protocol to accommodate low-input analysis of RNAs up to 120 nucleotides in size. To our knowledge, our study is the first to provide the complete length distribution profile of both type I and the longer type II tRNAs, typically ranging from 75-90 nucleotides in size^63^ in EVs, both *in vitro* and *in vivo*. In contrast to earlier observations^19,64,65,35^, we detected only a minimal fraction of fragmented tRNAs associated with EVs. The majority of tRNA reads are full-length sequences without an intact 3’ CCA tail, which was confirmed with Northern blotting (Figure 1F). In agreement with a previous report^47^, full-length tRNAs are near-absent in the NVEP fractions, which are enriched in fragmented ncRNAs primarily tRNA halves. Notably, the design of our study does not allow discrimination between ‘true’ tRNA halves or folded nicked tRNAs, as shown in previous studies^34,64^.

Proteinase K-RNase I treatment of our EVs yielded near identical results indicating full-length tRNAs minus an intact 3’ CCA are present within the lumen of EVs. In addition, the characterization of misincorporation signatures and truncated 3’ ends in the present study revealed that EV-tRNAs were, in fact, mature tRNA molecules that had apparently undergone complete processing prior to their sorting into the lumen of EVs. While the majority of identified modification-induced mismatches in EV-associated tRNA read sequences corresponded to their cellular counterparts, we noted differences in guanine modifications at position 26. This mismatch position aligns well with the m^2^ G modification that is widespread in tRNAs. Importantly, this misincorporation signature was previously detected and verified with sequencing-based methods using the ultra-processive group II intron maturase, TGIRT^57^, and is known to enhance stability of the coaxially stacked D and anticodon stems^66,67^. Notably, base-methylated G-derivatives achieve their maximum modification density after 48 hours^68^, providing further evidence that EV-tRNAs likely originated from the fully mature and long-lived cytoplasmic tRNA population prior to sorting into EVs. Nevertheless, it remains to be determined whether these tRNA modifications play a role in the preferential sorting into EVs.

The presence of entirely processed, full-length tRNAs inside EVs raises the question into the function of their inclusion as cargo inside the lumen of EV preparations. In the cell, tRNAs play a pivotal role as central adapter molecules in translating the genetic code into proteins^69^. The full set of 20 amino acids are encoded by a total of 61 possible codons, which are read by tRNA isoacceptors that exhibit significant variation in their cellular concentrations^70^. The levels of RNAs in the cell are mainly regulated via a dynamic balance between their synthesis rate (transcription in the nucleus, processing/maturation) and their degradation in the cell’s cytoplasm, thereby controlling the availability of RNA for various cellular processes^71^. Alongside tRNA concentrations^72^, factors such as variation in aminoacylation^73^, CCA tail integrity^74^, and the types and abundance of their modifications also influence protein translation efficiencies. Previous studies suggested that EV-sorted tRNAs as well as circulating tRNAs in human plasma harbored posttranscriptional modifications expected for mature tRNAs, and in most cases ended with the post-transcriptionally added 3′ CCA, although many truncated species were detected^33^. We demonstrate here that 3’ CCA tail shortening is pervasive across all mature tRNA types inside bulk EV preparations. Remarkably, we showed that at times the tRNA 3’ CCA tail shortening extends several nucleotides into the tRNA, suggestive of exonuclease cleavage, which was not observed for the tRNA 5’ ends. This observation raises the intriguing possibility that exporting completely processed, functional mature full-length tRNAs into EVs is actually rare and that sorting into EVs is a mechanism to maintain a fully functional cytoplasmic tRNA pool. While cells efficiently add and repair CCA ends via the CCA-adding enzyme TRNT1 to enhance the number of intact tRNAs during high demand for protein synthesis, under conditions of ample supply, excess highly stable tRNAs^75^ could potentially get encapsulated and shuttled out of the cell. Because most tRNAs in a cell are maintained in an aminoacylated state, removing the CCA, prior incorporating tRNAs in EVs would ensure that amino acid building blocks, which are also used to power metabolism, particularly in cancer cells, would remain available in the cell^76^.

Yip et al.^77,78^ suggested an additional regulatory role for the CCA ends during translation. They linked the 3’ CCA removal to ribosome-associated quality control, which is triggered by aberrantly stalled translation complexes. The faulty peptidyl-tRNAs on stalled ribosomes are specifically processed for recycling or degradation to prevent further use of non-functional tRNAs in translation. Hence, sorting faulty tRNAs into EVs could also constitute an alternative mechanism to alleviate this cellular stress. It is worth noting that the 3’ CCA ends of tRNAs have also been implicated in several other regulatory pathways^61^, such as the sequential deactivation at the 3’ CCA end of tRNAs in response to stresses, leading to reversible translational repression at very low metabolic costs^74,79^. However, the exact molecular mechanism responsible for the absence of 3’ CCA tails within EVs remains unknown and will be the subject of our future investigations.

While we successfully reduced biases caused by secondary structures and posttranscriptional modifications, certain biases remained unaddressed in the present study. Consequently, we might not have captured all transcripts, like fragmented RNAs and nicked tRNAs, that may contain a 2’-3’ cyclic phosphate at the 3’ ends of the RNA, for instance. Additionally, the size range of EVs isolated from biofluids overlaps with other particles of similar dimensions which could have partially interfered with our analysis^80^. Furthermore, considering the diverse cellular origins of EVs in circulation, the resulting signal could display significant heterogeneity. Nonetheless, our analysis of pEVs revealed comparable class and size distributions, as well as nucleotide misincorporation signatures, in comparison to results obtained from cultured cell-derived EVs. Lastly, further tailoring of the library preparation conditions for quantification of EV-small ncRNAs in the size range of 50-120 nt could benefit from, for instance, incorporation of unique molecular identifiers (UMIs) to reduce PCR amplification bias of unwanted smaller sequences. Hence, targeted sequencing of either the general population or chemically modified small ncRNAs in the size range of 18-50 nt, or the larger range of 50-120 nt, could further improve the accurate quantification of small ncRNAs in low-input samples. Finally, we cannot exclude the possibility that the tRNAs we measured in our bulk EV preparations are associated with a specific subtype of EVs. EV subtypes include exosomes, which are 30-150nm sEVs that may originate from multivesicular bodies (MVBs) as intraluminal vesicles (ILVs) released upon MVB-plasma membrane fusion. However, sEVs can also bud directly from the plasma membrane and may be biochemically similar to ILV-derived exosomes. Conversely, micro-vesicles (MVs) are categorized as lEVs with a diameter ranging from 150 to 1000 nm that also bud directly from plasma membrane^83^. This makes the biochemical differentiation between MVs and sEVs not always straightforward^84^. At least in the case of our EV preparations derived from *in vitro* cultured cell EVs, that likely contained small and large EVs of various subcellular origin, we favor the hypothesis that the majority of EV-associated tRNAs, regardless of the EV subtypes, lacks an intact 3’ CCA. Future studies will be required to decipher whether specific EV subtypes are enriched in full-length tRNAs and whether different subtypes harbor tRNAs with and without complete CCA-tails. In addition, EVs used in this work were purified from the supernatant of cells propagated under standard culture conditions, while pEVs are probably derived from many different cell types in various cell states. Nevertheless, it is conceivable that the tRNA content and integrity could be modulated in response to activation of signaling pathways, as suggested by prior studies^35,85^.

In summary, our data revealed a predominance of full-length ncRNAs in bulk EVs derived from cell-culture supernatants as well as circulating plasma EV preparations of diverse tissue origin. Our results strongly suggest that the tRNAs located in the lumen of EVs, although fully processed and likely originating from mature molecules, harbor a truncated 3’ CCA tail rendering them incompetent for mRNA translation. Further investigations aimed at understanding the role for extracellular tRNAs could shed light on potential strategies used by cells to control protein synthesis and tRNA demand during physiological stress conditions.

## DATA AND CODE AVAILABILITY

The ALL-tRNAseq computational pipeline is available at https://github.com/bioinfoUGR/sRNAtoolbox.

## MATERIAL AND METHODS

### Cell culture

MDA-MB-231 and HEK293T cells were cultured at 37°C in a humidified atmosphere containing 5% CO2 in Dulbecco’s Modified Eagle Medium (DMEM, Thermo Fisher Scientific, cat# 41965039) with 1x MEM non-essential amino acids solution (ThermoFisher Scientific, cat# 11140-035) for media used with HEK293T cells. HCT116 cells were cultured in McCoy’s 5A medium (Lonza, cat# 12-688F). All passaging of adherent cell lines was performed with trypsin-ethylenediaminetetraacetic acid (EDTA) solution (Thermo Fisher Scientific, cat# 15400054) according to the manufacturer’s instructions. Lymphoblastoid RN cells were cultured in suspension at 37°C with 5% CO2 in Roswell Park Memorial Institute (RPMI)-1640 medium with Hepes (Thermo Fisher Scientific, cat# 22400089). All media were supplemented with 10% heat-inactivated fetal bovine serum (FBS; LPS, cat# S-001A-BR) and and 100U/ml penicillin G and 100 μg/ml streptomycin. All cell lines were subjected to mycoplasma testing and only used for experiments when confirmed negative.

### Blood collection and plasma preparation

All blood samples that were used in this study were collected longitudinally from four classic Hodgkin lymphoma (cHL) patients, at two different timepoints through biobanking and were obtained under the approval of the Medical Ethical Committee of Amsterdam UMC. Biobank is registered under 2018.359. Sample collection procedures were conducted in accordance with the Declaration of Helsinki and Good Clinical Practice guidelines. Complete data on clinical features at presentation, treatment and outcome as well as pathology reports were available for all samples.

Blood samples were collected using plasma collection tubes (BD Vacutainer, Ethylene Diamine Tetra Acetic Acid (EDTA)) and were processed within 2 hours after collection. Tubes were processed through two rounds of centrifugation at room temperature, initially at 900 g for 7 minutes, followed by a second centrifugation at 2500g for 10 minutes. All plasma samples were collected as 1 ml aliquots and stored at −80C until further use, and were not subjected to any freeze-thaw cycles prior to analysis.

### Sample fractionation

For all cell lines, culture supernatants were collected and underwent sequential centrifugation twice at 500g for 10 minutes each, and 2x 2000g for 15 minutes to remove dead cells and cellular debris. EVs, NVEP fractions and plasma-derived extracellular vesicles (pEVs) were subsequently isolated through Size Exclusion Chromatography (SEC), following the procedure previously described by Eijndhoven et al^43^ with the following modifications. Cell line supernatants were first concentrated via filtration using a Centricon Plus-70 Centrifugal Filter with a 100kDA cut-off (Millipore, cat# UFC710008). EV and NVEP fractions as well as pEVs were then collected using an Automatic Fraction Collector (AFC, IZON Science LTD.) with qEV original 70nm columns (IZON Science LTD., cat# SP1) with a void volume of 2.85 ml. EVs of interest were obtained in 500 μl volumes from fractions 3 and 4, while SEC NVEP fractions of interest were collected in 500 μl volumes from fractions 14 and 15. We determined the enrichment of EVs and NVEPs by western blotting, electron microscopy (EM) and particle measurements, according to MISEV guidelines. Subsequently, each fraction was subdivided into 250 μl aliquots, followed by a 15-minute incubation at room temperature with 1.25 ml Qiazol lysis reagent (Qiagen, cat# 79306). All samples were stored at 80°C until further processing.

### RNAse I sensitivity assay

EVs were treated with 50 µg/ml Proteinase K for 30 minutes on ice (Sigma cat# P2308), followed by inactivation with 25 mM PMSF (Sigma cat# P7626). Subsequently, the samples were treated with 500 units RNase If for 20 minutes at 30°C (NEB cat# M0243S).

### RNA isolation

Total RNA of cell lines was extracted using miRNeasy micro Kit (Qiagen, cat# 217084) according the manufacturer’s protocol. RNA from EVs, SEC NVEP fractions, and pEVs were isolated using miRNeasy serum/plasma kit (Qiagen, cat# 217184). Total RNA isolation was performed in an RNase-free environment. The complete EV fractions, NVEP fractions and pEV fractions were isolated using one miRNeasy column each and all samples were eluted in 14 μl H_2_O. tRNA deacylation was then performed on all samples by incubating 2 μg of cellular total RNA or a volume of 4 μl of extracellular total RNA in 0.1 M Tris-HCL pH9.0 and 1 mM EDTA for 30 minutes at 37°C. After ethanol precipitation in presence of 15 μg GlycoBlue Coprecipitant (Invitrogen, cat# AM9516), total RNA was resuspended in RNase-free H_2_O.

Quality and integrity of all cellular RNA samples were measured on a NanodropOne (ThermoFisher Scientific) and Agilent 2100 Bioanalyzer (Agilent Technologies), using an RNA 6000 Nano kit (Agilent Technologies, cat# 5067-1511). Quality and integrity of extracellular RNA was evaluated by quality control PCR with a 1:10 diluted sample. The input for the QC-PCR reaction was 3 μl of 1:10 diluted RNA that was reverse transcribed using TaqMan^™^ MicroRNA Reverse Transcription kit (ThermoFisher Scientific, cat# 4366596). For EVs from culture supernatants, the multiplex reaction contained primers for hsa-miR-21-5p and hsa-let7a-5p. For pEVs, the multiplex reaction contained primers for hsa-miR-486-5p, hsa-miR-21-5p, hsa-miR10b-5p, and hsa-let7a-5p, as described by Eijndhoven et al^86^. Nuclease-free water was added to the reaction to a volume of 50 μl and subsequently 3 μl of cDNA was PCR-amplified in 40 cycles of 95°C for 15 seconds and 60°C for 1 minute on an ABI 7500 Fast System. All samples were measured in duplicates and data were analyzed using 7500 Software v2.0.6.

### Demethylation reaction

Demethylation reactions were adapted from previously published methods^87–89^. A total of 2 μg of cellular total deacylated RNA or a volume of 4 μl extracellular total deacylated RNA for each cell line and plasma sample underwent treatment with 19 μM of recombinant *E. Coli* AlkB, comprised of 8.5μM AlkB WT and 10.5 μM AlkB-D135S mutant. The reaction mixture also included 10 mM KCl, 2 mM MgCl_2_, 283 μM freshly prepared (NH_4_)_2_Fe(SO_4_)_2·_ 6H_2_O, 0,3 mM 2-ketoglutarate, 2 mM freshly made L-ascorbic acid, 40U RNase Inhibitor and 50 mM MES buffer pH 5.0 (Fisher Scientific, cat# 15474529), all in a total volume of 40 μl. Incubation of reactions occurred at 25°C for a duration of 2h and was subsequently quenched with 5 mM EDTA (Ambion, cat# AM9260). RNA was then recovered using an Oligo Clean&Concentrator-5 kit (ZYMO Research, cat# D4060) and eluted in 6 μl nuclease-free water (Ambion, cat# AM9937).

### Small RNA-seq library preparation

Sequencing libraries were prepared following the ALL-tRNAseq procedure described by Scheepbouwer et al^31^, with the following modifications. Briefly, a 5’ phosphorylated 3’ end adapter featuring 4 randomized nucleotides at the 5’ terminus and a 3’ blocking group (3C Spacer; 3SpC3, IDT) underwent adenylation using Mth RNA ligase (New England Biolabs, cat# M2611A). Adapters were subsequently ligated to deacylated and demethylated RNA templates using a truncated KQ T4 RNA ligase 2 (New England Biolabs, cat# M0373L) for 1 hour at 25°C with 20% PEG8000 (New England Biolabs). After ligation, RNA was gel-purified and subjected to size-fractionation on a denaturing polyacrylamide gel (Novex TBE-urea 10%, Invitrogen, cat# EC68752BOX)) to enrich for small RNA molecules. Two FAM-markers of 25 nt and 39 nt, along with the low range ssRNA ladder (New England Biolabs, cat# N0364S), were used to guide size-fractionation. Gels were briefly washed and stained using 1x SYBR gold nucleic acid stain (Thermo Fisher Scientific, cat# S11494) in 1x TBE for 10 minutes, and 3’adapter-ligated-RNA was excised on a blue light transilluminator with a size range of 50-140 nt for cell line-derived cells and EVs, and 39-140 nt for the SEC NVEP fractions and plasma-derived EVs. RNA recovery from the excised polyacrylamide gel pieces was performed by crushing the gel fragments and soaking them into 0.3M NaCl overnight at 4°C, followed by ethanol precipitation. Subsequently, 5’ adapters with 4 terminal randomized nucleotides at the 3’ end was added using T4 RNA ligase (New England Biolabs, cat# M0204S). Ligation was carried out for 1 hour at 25°C with 20% PEG8000. After ligation, reverse transcription was performed for 1h at 42°C using MarathonRT (Kerafast, cat# EYU007). Reactions were terminated by incubation with 0.25 M NaOH, at 95°C for 3 min, followed by neutralization with 0.25 M HCl. The ensuing cDNA was purified using the MinElute Reaction Cleanup kit (Qiagen, cat# 28006) and PCR amplified using Phusion high-fidelity PCR master mix (Fisher Scientific, cat# F531L) for 12 cycles of 98°C for 5 s, 62°C for 10s and 72°C for 10 s. An additional size-fractionation step was performed for all EV and NVEP fractions using a Novex TBE PAGE gel (6%, Invitrogen, Cat# EC6265BOX) with a size range of 150 – 400 bp for cell line-derived cells and EVs, and 140 - 400 bp for the SEC NVEP fractions and plasma-derived EVs. Size determination employed the molecular marker Quick-load pBR322 DNA-Mspl Digest, designed for small RNA library preparations (New England Biolabs, cat# N3032S). Primers, randomized adapters, Illumina Multiplex and barcode primers were obtained from IDT and are listed in Supplemental Table 1. Quality control for the sequencing libraries involved the use of an Agilent high sensitivity DNA kit (Agilent Technologies) and Fragment Analyzer Systems (Agilent Technologies) to assess size, purity and concentration. Subsequently, the ALL-tRNAseq small RNA libraries underwent single-end sequencing for 150 cycles on NovaSeq 6000 at GenomeScan B.V. Leiden, Netherlands.

### Sequencing Analysis

Sequencing analysis was performed as described by Scheepbouwer et al.^31^. Briefly, quality control and adapter sequence trimming were performed using sRNAbench^90^. For all small RNA classes, a first round of read mapping was performed using the standard sRNABench pipeline via bowtie. A second round of alignment using a Smith–Waterman BioJava implementation was used to retrieve additional tRNA reads containing gaps and sequence mismatches. Reference libraries from GtRNAdb 2.0 (Homo sapiens, GRCh38/hg38) were utilized for nuclear tRNAs, while sequences from mitotRNAdb were used for mitochondrial tRNAs. tRNA expression was examined at multiple levels, including tRNA gene level and anticodon level. Only tRNA reads containing an intact 5’ end with at least partial *CCA* ending were classified as mature full-length transcripts.

### Quantitative PCR (qPCR) for CCA-tail analysis

NEBNext 3’ SR adaptors for Illumina (New England Biolabs, cat# E7332) were ligated to the 3’ ends of total RNA from EVs and cellular samples isolated from an RN cell line using a truncated KQ T4 RNA ligase 2 (New England Biolabs, cat# M0373L) for 1 h at 25°C. Ligated RNA was subsequently reverse transcribed in a 20 ul reactions using SuperScript IV (ThermoFisher Scientific, cat(# 18090010) for 10 minutes at room temperature (RT), 25 minutes at 50°C and reactions were terminated by incubation for 10 minutes at 80°C. Reaction volumes were increased to 50 ul and 3 μl was subsequently used for qPCR reactions with SYBR Green (Roche, cat# 04887352001). Forward primers were designed at the 5’ ending of tRNAs and the reverse primers were designed against the 3’adapter while specificity was created against the 3’ endings. PCR amplification was performed for 45 cycles of 95°C for 10s, 60°C for 15s and 72°C for 15s. Cycle threshold (Ct) values were obtained and ΔΔCt values were calculated to determine relative abundance and reported as mean ±SD. Primer sequences are available in Supplemental Table 1.

### Statistical analysis

All expression values derived from sequencing data presented throughout this work were expressed as RPM (Reads Per Million) per reference library or total mapped reads, as indicated in Figure legends. The webserver NormSeq^91^ was used for best data normalization based on information gain analysis. Differential expression analysis was performed with the EdgeR^92^ package on the Galaxy web platform^93^ (Galaxy Tool Version 3.36.0) using Relative Log Expression (RLE) normalization for all samples. Count files were uploaded, applying standard filtering except filtering of lowly expressed tRNA anticodons below 100 counts. Benjamini-Hochberg’s FDR is applied to the p values in order to correct for multiple testing. Statistical analyses were performed using GraphPad Prism9 Software. Significance for modification analyses was calculated using Student’s t-tests. Pearson correlations were calculated using the function pearsonr from the scipy package (v1.8.1).

### Northern Blotting

Northern blots were largely performed as initially described by Gerber et al.^94^, with additional modifications. A 4 μl total RNA sample was separated on a 10% urea-polyacrylamide gel. After electrophoresis, RNAs were transferred to a HYBOND N+ membrane (Thermofisher, cat# RPN2020B) using a wet electrophoretic transfer system at 30V for 1 hour. Following transfer, the RNAs were cross-linked with UV light (0.24J on the RNA side and 0.12J on the opposite side of the membrane). Pre-hybridization was performed in ULTRAhyb Oligo Hybridization buffer (Thermofisher, cat# AM8663) with gentle agitation for 1 hr at 42°C. Hybridization then occurred overnight at 37°C with 50 nM biotinylated oligonucleotide probes. After hybridization, the membranes underwent two 10-minute washes with 5X SSC, 0.5% SDS at room temperature. Subsequently, membranes were blocked for 15 minutes at room temperature using 2X SSC, 0.5% SDS and 3% BSA. Membranes were then incubated with Streptavidin-HRP (1:40,000, Thermofisher N100) in 2X SSC containing 3% BSA and 0.5% SDS for 30 minutes. Membranes were washed with ABS buffer (10% BSA and 1% Triton X-100 in 2X SSC) and subjected to two 5-minute washes with 2X SSC. Before exposure, membranes were briefly rinsed with PBS and then subjected to detection using SuperSignal WestPico Plus ECL-based detection (ThermoFisher, cat# 34580). Finally, membranes were exposed on a Uvitec chemiluminescence imaging system. Oligonucleotide sequences are provided in Supplemental Table 1.

## ACKNOWLEDGEMENTS

This work was supported by funding awarded to D.M.P. provided by the Stichting CCA [CCA2021-9-77; CCA2023-9-93], MRD Hodgkin Lymphoma, and the AQrate TKI-project from Health-Holland. Work in the lab of A.G. was supported by the NWO Talent Programme Vidi grant [VI.Vidi.193.107] and Stichting Cancer Center Amsterdam [CCA2021-5-26]. E.A. was supported by the National Institute of General Medical Sciences [R01GM139928]. C.G. was supported by the Stichting Cancer Center Amsterdam [CCA2021-9-77; CCA2023-9-93]. The authors would like to acknowledge the usage of the computational infrastructure of the Computational Epigenomics Lab of the University of Granada.

---

**Supplemental Figure 1.** (A) RNA class distribution in percentage of total normalized reads for the ALL-tRNA-seq library preparation protocol in the EV fraction of two cell lines of different origin (RN, MDA-MB-231). (B-D) Pearson correlation between normalized full-length tRNA reads in cells and normalized full-length tRNA reads in EVs of (B) HCT116 (n=1; *r*: 0.78), (C) MDA-MB-231 (n=1; *r*: 0.83), and (D) 293T (n=1; *r*: 0.84). (E) SYBR gold staining of untreated tRNAs (left lane) and upon treatment with RNase I (right lane), isolated from U118 cells. Ladder size is indicated next to the image. (F) Proportion of nucleotide variations matching to acp^3^U20, m^2^2G26, I34, and m^1^I37 in tRNA-AlaAGC from cells (n=3; blue) and EVs (n=3; red). Significance was calculated using a Student’s t-test. (G) Comparison between cells and EVs in the proportion of tRNA reads containing the tRNA 5’ end. (H) Evaluation of tRNA-GluCTC CCA 3’ ending in cells and EVs using qPCR. Ct values are indicated on the Y axis. RNY1 was included as a control. (I) SYBR gold staining of qPCR products of tRNA-GluCTC with and without CCA 3’ ends. (J) Violin plot showing read length distributions in percentage of Y RNA from 4 cell lines of different origin (RN, MDA-MB-231, HCT116, and HEK293T).

