## Supplementary figures and images for "Full-length tRNAs lacking a functional CCA tail are selectively sorted into the lumen of extracellular vesicles"

### Supplemental Figure 1

**Supplemental Figure 1.**

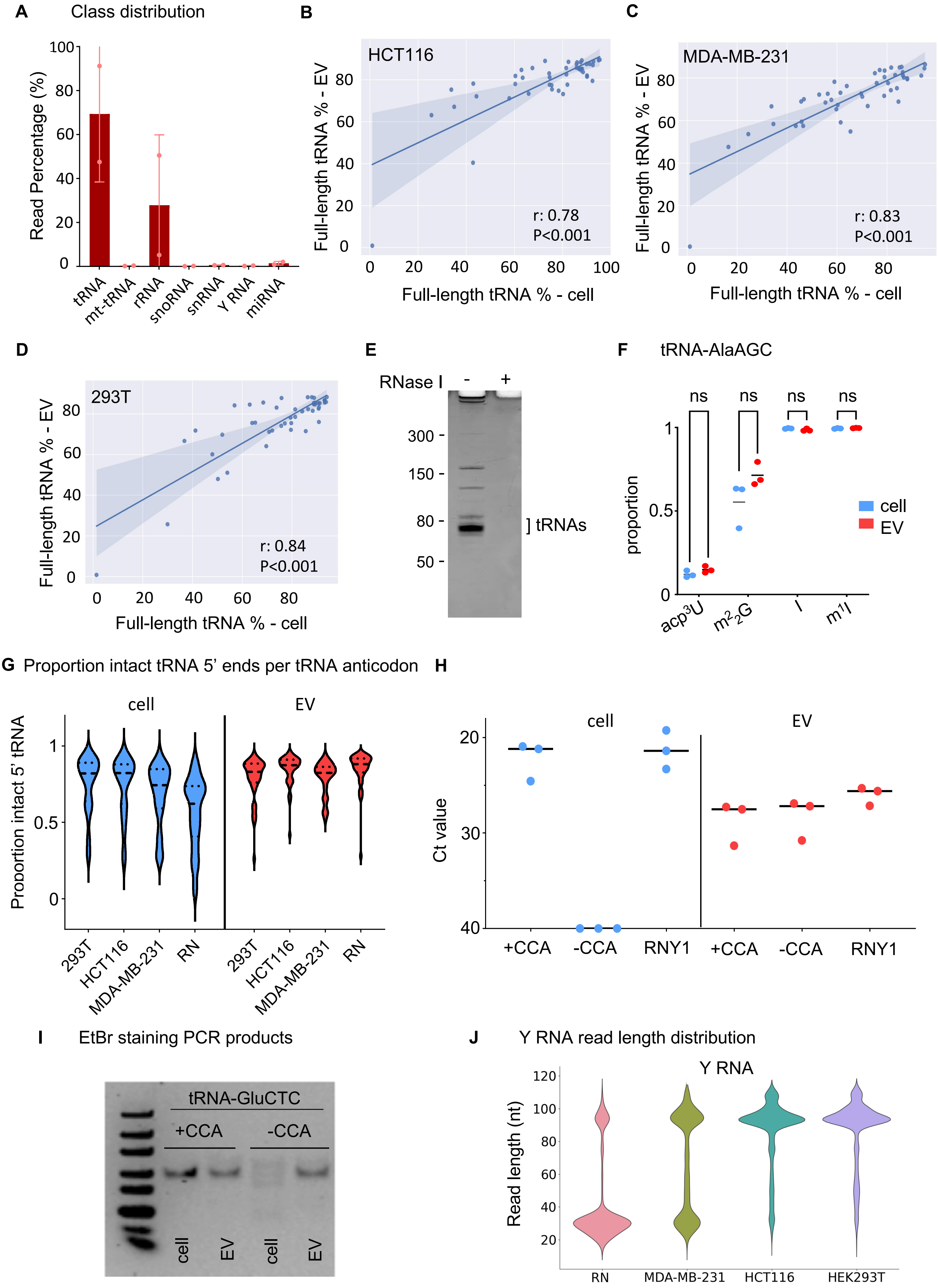
