## Supplemental Table 1 for "Full-length tRNAs lacking a functional CCA tail are selectively sorted into the lumen of extracellular vesicles"

| Primers for qPCR and library construction, oligonucleotides and probes for Northern Blotting | |
|---|---|
| tRNA-GluCTC FW | 5'- TCCCTGGTGGTCTAGTGGTT -3' |
| tRNA-GluCTC CCA RV | 5'-CGTGTGCTCTTCCGATCTTGG -3' |
| tRNA-GluCTC CCAless RV | 5'- CGTGTGCTCTTCCGATCTTTC -3' |
| 3´ SR Adaptor for Illumina | 5’-AGATCGGAAGAGCACACGTCT/3SpC3 |
| degenerate 3´ SR Adaptor for Illumina-4N | 5´/5Phos/(N:25252525)(N)(N)(N)(N)AGATCGGAAGAGCACACGTCT/3SpC3 |
| degenerate 5´ SR Adaptor for Illumina-4N | 5´-rGrUrUrCrArGrArGrUrUrCrUrArCrArGrUrCrCrGrArCrGrArUrCr(N:25252525)r(N)r(N)r(N)r(N)-3´ |
| 23nt-RNA marker-FAM | 5'-/6-FAM/rUrGrUrArCrGrCrGrArArUrArGrUrUrArArArCrCrUrG/ddC-3' |
| 39nt-RNA marker-FAM | 5'-/6-FAM/rUrUrArGrArArGrArCrGrArGrUrCrArCrCrArUrGrArCrCrArArUrArGrCrGrArUrArArCrUrGrA/ddC-3' |
| NEBNext SR RT Primer for Illumina | 5´-AGACGTGTGCTCTTCCGATCT-3´ |
| SR Primer for Illumina (RP1) | 5´-AATGATACGGCGACCACCGAGATCTACACGTTCAGAGTTCTACAGTCCGA-3´ |
| RPI02 PCR index primer | 5´-CAAGCAGAAGACGGCATACGAGATACATCGGTGACTGGAGTTCAGACGTGTGCTCTTCCGATC*T-3´ |
| RPI03 PCR index primer | 5´-CAAGCAGAAGACGGCATACGAGATGCCTAAGTGACTGGAGTTCAGACGTGTGCTCTTCCGATC*T-3´ |
| RPI07 PCR index primer | 5´-CAAGCAGAAGACGGCATACGAGATGATCTGGTGACTGGAGTTCAGACGTGTGCTCTTCCGATC*T-3´ |
| RPI09 PCR index primer | 5´-CAAGCAGAAGACGGCATACGAGATCTGATCGTGACTGGAGTTCAGACGTGTGCTCTTCCGATC*T-3´ |
| RPI11 PCR index primer | 5´-CAAGCAGAAGACGGCATACGAGATGTAGCCGTGACTGGAGTTCAGACGTGTGCTCTTCCGATC*T-3´ |
| RPI12 PCR index primer | 5'-CAAGCAGAAGACGGCATACGAGATTACAAGGTGACTGGAGTTCAGACGTGTGCTCTTCCGATC*T-3´ |
| RPI17 PCR index primer | 5´-CAAGCAGAAGACGGCATACGAGATCTCTACGTGACTGGAGTTCAGACGTGTGCTCTTCCGATC*T-3´ |
| RPI18 PCR index primer | 5´-CAAGCAGAAGACGGCATACGAGATGTGCGGACGTGACTGGAGTTCAGACGTGTGCTCTTCCGATC*T-3´ |
| RPI21 PCR index primer | 5´-CAAGCAGAAGACGGCATACGAGATCGAAACGTGACTGGAGTTCAGACGTGTGCTCTTCCGATC*T-3´ |
| RPI22 PCR index primer | 5´-CAAGCAGAAGACGGCATACGAGATCGTACGGT GACTGGAGTTCAGACGTGTGCTCTTCCGATC*T-3´ |
| RPI23 PCR index primer | 5´-CAAGCAGAAGACGGCATACGAGATCCACTCGTGACTGGAGTTCAGACGTGTGCTCTTCCGATC*T-3´ |
| RPI24 PCR index primer | 5´-CAAGCAGAAGACGGCATACGAGATGCTACCGTGACTGGAGTTCAGACGTGTGCTCTTCCGATC*T-3´ |
| RPI25 PCR index primer | 5'-CAAGCAGAAGACGGCATACGAGATATCAGTGTGACTGGAGTTCAGACGTGTGCTCTTCCGATC*T-3' |
| RPI26 PCR index primer | 5'-CAAGCAGAAGACGGCATACGAGATGCTCATGTGACTGGAGTTCAGACGTGTGCTCTTCCGATC*T-3' |
| RPI27 PCR index primer | 5'-CAAGCAGAAGACGGCATACGAGATAGGAATGTGACTGGAGTTCAGACGTGTGCTCTTCCGATC*T-3' |
| RPI28 PCR index primer | 5'-CAAGCAGAAGACGGCATACGAGATCTTTTGGTGACTGGAGTTCAGACGTGTGCTCTTCCGATC*T-3' |
| RPI29 PCR index primer | 5'-CAAGCAGAAGACGGCATACGAGATTAGTTGGTGACTGGAGTTCAGACGTGTGCTCTTCCGATC*T-3' |
| RPI30 PCR index primer | 5'-CAAGCAGAAGACGGCATACGAGATCCGGTGGTGACTGGAGTTCAGACGTGTGCTCTTCCGATC*T-3' |
| RPI31 PCR index primer | 5'-CAAGCAGAAGACGGCATACGAGATATCGTGGTGACTGGAGTTCAGACGTGTGCTCTTCCGATC*T-3' |
| RPI32 PCR index primer | 5´-CAAGCAGAAGACGGCATACGAGATTGAGTGGTGACTGGAGTTCAGACGTGTGCTCTTCCGATC*T-3´ |
| RPI33 PCR index primer | 5´- CAAGCAGAAGACGGCATACGAGATCGCCTGGTGACTGGAGTTCAGACGTGTGCTCTTCCGATC*T-3´ |
| RPI34 PCR index primer | 5'-CAAGCAGAAGACGGCATACGAGATGCCATGGTGACTGGAGTTCAGACGTGTGCTCTTCCGATC*T-3' |
| RPI35 PCR index primer | 5'-CAAGCAGAAGACGGCATACGAGATAAAATGGTGACTGGAGTTCAGACGTGTGCTCTTCCGATC*T-3' |
| 5'GlyGCC-5-1-northern probe | /5Biosg/TCTACCACTGAACCACCAAT-3' |
